# *In vitro* synthesis and reconstitution using mammalian cell-free lysates enables the systematic study of the regulation of LINC complex assembly

**DOI:** 10.1101/2021.04.11.439350

**Authors:** Sagardip Majumder, Yen-Yu Hsu, Hossein Moghimianavval, Michael Andreas, Tobias W. Giessen, G.W. Gant Luxton, Allen P. Liu

**Author notes:** Institute of Protein Design, University of Washington, Seattle, Washington, 98195, USA. Contributed equally (Yen-Yu Hsu, Hossein Moghimianavval). Corresponding author: Allen P. Liu, **Email:**. **Author Contributions:** S.M., G.W.G.L., and A.P.L. conceived the project and designed experiments. S.M., Y.Y.H., H.M. carried out the experiments and prepared figures. M.A. and T.W.G. designed and performed the cryo-EM experiment. S.M. analyzed relevant data. S.M., G.W.G.L., and A.P.L. wrote the manuscript.

## Abstract

Membrane proteins perform numerous important functions in cells and tissues. Approximately 20% of the human genome encodes for membrane proteins, which represent the majority of targets for clinically relevant small molecules. Consequently, understanding their structure and structure-function relationships is a fundamental problem in biomedical research. Given the difficulties inherent to performing mechanistic biochemical and biophysical studies of membrane proteins *in vitro*, we previously developed a facile HeLa cell-based cell-free expression (CFE) system that enables the efficient reconstitution of full-length (FL) functional membrane proteins in supported lipid bilayers. Despite having shown the directional reconstitution of CFE-synthesized FL inner nuclear membrane SUN proteins (i.e. SUN1 and SUN2), which directly interact with outer nuclear membrane KASH proteins within the nuclear envelope lumen to form linker of nucleoskeleton and cytoskeleton (LINC) complexes that mechanically couple the cytoskeleton and nucleus, the mechanism underlying regulated LINC complex assembly remains unclear. Here, we provide evidence that suggests that the reconstitution of CFE-synthesized FL membrane proteins in supported lipid bilayers occurs primarily through the fusion of endoplasmic reticulum-derived microsomes present within our CFE reactions with our supported lipid bilayers. In addition, we demonstrate the ease with which our synthetic biology platform can be used to investigate the impact of the chemical environment (e.g. calcium ions and redox state) on the ability of CFE-synthesized FL SUN proteins reconstituted in supported lipid bilayers to interact with the luminal domain of the KASH protein nesprin-2. Moreover, we use our platform to study the molecular requirements for the homo- and hetero-typic interactions that can occur between SUN1 and SUN2. Finally, we show that our platform can be used to simultaneously reconstitute three different CFE-synthesized FL membrane proteins in a single supported lipid bilayer. Overall, these results establish our HeLa cell-based CFE and supported lipid bilayer reconstitution platform as a powerful tool for performing mechanistic dissections of the oligomerization and function of FL membrane proteins *in vitro*. While our platform is not a substitute for cell-based studies of membrane protein biochemistry and function, it does provide important mechanistic insights into the biology of difficult-to-study membrane proteins.

**ABSTRACT GRAPHIC:** 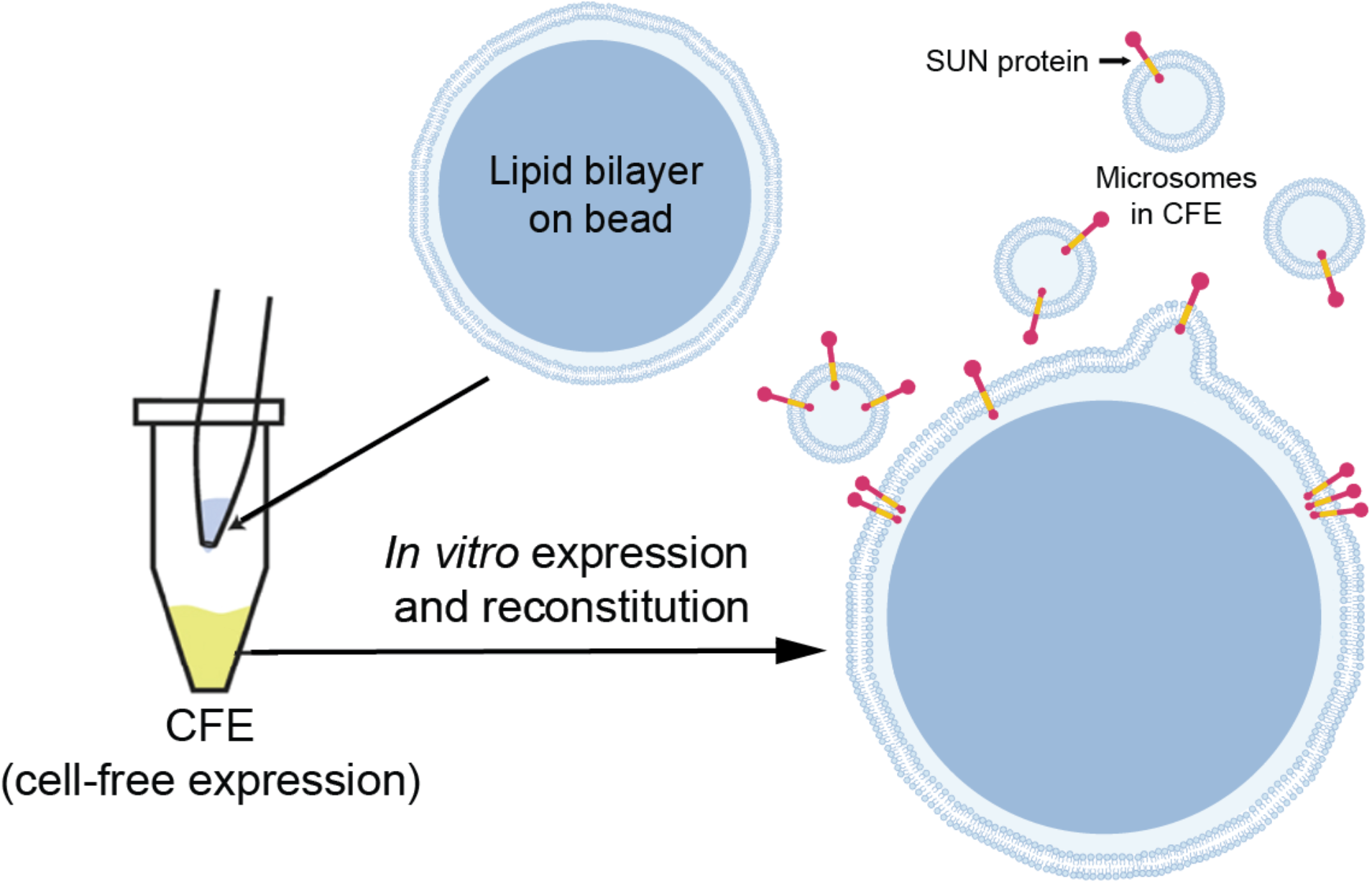

## INTRODUCTION

A fundamental aspect of eukaryotic cellular biology is the establishment of membrane-enclosed organellar compartments, such as the endoplasmic reticulum (ER) and the nuclear envelope (NE). These compartments are critically important for the physical segregation of organelle-specific chemical reactions and signal transduction cascades as well as for their movement and positioning within cells. Integral and/or peripheral membrane proteins mediate many of these functions and their importance is highlighted by the fact that membrane proteins are encoded by ~20% of the genes present within human genome^1^. In addition, the majority of therapeutics target membrane proteins^2^, which represent > 60% of all United States Food and Drug Administration-approved small molecule drugs^3^. Despite their status as clinically relevant therapeutic targets, we know very little about the biochemistry, biophysics, or structural biology of FL membrane proteins. For example, < 1% of all determined protein structures belong to membrane protein families^4^.

Several practical experimental reasons exist to explain this dearth of information regarding FL membrane proteins. Because of their nature, membrane proteins possess hydrophobic surfaces, and they can only be extracted from cell membranes with detergents. Moreover, experimentally isolated membrane proteins are notoriously unstable. Consequently, working with FL membrane proteins is challenging at all levels, including their expression, solubilization, purification, and crystallization^5^. To begin to address these challenges, we previously described the development of a HeLa cell-based cell-free expression (CFE) system that can be used as a simple “bottom-up” synthetic biology platform^6,7^ for the rapid reconstitution of FL inner nuclear membrane (INM) Sad1/UNC-84 (SUN) proteins in supported lipid bilayers, which we called artificial nuclear membranes (ANMs)^8^.

We originally developed ANMs to be able to reconstitute and mechanistically dissect the assembly of linker of nucleoskeleton and cytoskeleton (LINC) complexes, which are conserved NE-spanning molecular bridges that mediate several fundamental cellular processes, including nuclear integrity, migration, and mechanotransduction^9–11^. LINC complexes are formed by the direct transluminal interactions of SUN proteins with the outer nuclear membrane (ONM)-localized Klarsicht/ANC-1/SYNE homology (KASH) proteins^12^ (**Fig. 1A**). *In vitro* studies performed using the purified C-terminal luminal domains (LDs) of KASH (e.g. the KASH peptide) and SUN proteins revealed that LINC complex assembly depends on the ability of SUN proteins to homo-trimerize, which results in the formation of KASH peptide-binding sites at the interaction interfaces between SUN protein protomers^13,14^.

**Figure 1.**
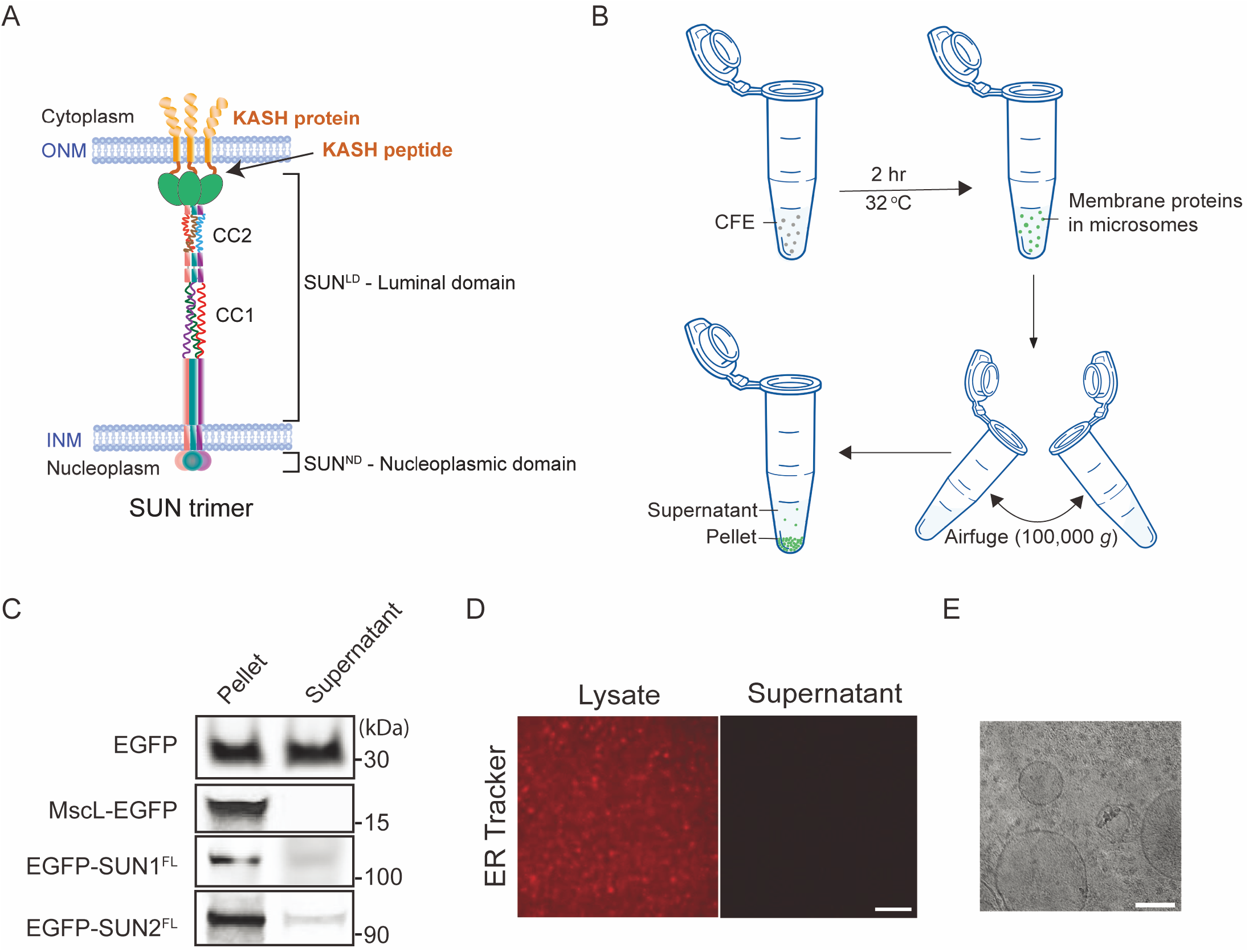
Microsomal structures and association of SUN proteins with membrane fragments in HeLa cell lysates used for CFE. A) Illustration of a SUN protein homo-trimer embedded in the INM that is interacting with KASH proteins embedded in the ONM. B) Schematic illustrating the use of Airfuge to spin down microsomal membranes in bulk CFE reactions. C) Representative in-gel fluorescence images of EGFP and the indicated EGFP-tagged proteins in supernatant and pellet fractions post ultracentrifugation. A self-inserting mechanosensitive membrane channel MscL was also expressed and served as a positive control. D) Representative confocal fluorescence images of ER tracker labeled HeLa lysate before and after spinning down microsomal membranes. Scale bar: 1 μm. E) Cryo-EM image of HeLa CFE system expressing EGFP-SUN1^FL^ showing vesicular structures. Small structures are visible in the background indicating ribosomes (~ 30 nm). Scale bar: 100 nm.

SUN1 and SUN2 are the most ubiquitously expressed mammalian SUN proteins^15^. We previously used ANMs to reconstitute CFE-synthesized N-terminally tagged FL mouse SUN1 and SUN2 (EGFP-SUN1^FL^ and EGFP-SUN2^FL^) in supported lipid bilayers^14^. EGFP-SUN1^FL^ and EGFP-SUN2^FL^ were expressed in the presence of supported lipid bilayers with excess membrane reservoirs (SUPER) templates to mediate their spontaneous reconstitution^16^. SUPER templates are formed on silica beads, which provide a suitable substrate for the incorporation of membrane-spanning proteins with the added advantage of their ease of handling and imaging. By carrying out protease protection assays on ANMs containing reconstituted CFE-synthesized EGFP-SUN1^FL^ or EGFP-SUN2^FL^, we found evidence for their directional insertion with their LDs facing the solvent accessible side of the SUPER templates. Based on the association of different truncated SUN protein constructs with SUPER templates, we also were able to predict that SUN2 possesses a single transmembrane domain and a hydrophobic membrane-associating domain, while SUN1 possesses three transmembrane domains. Furthermore, we demonstrated our ability to partially reconstitute LINC complex assembly through the interaction of reconstituted EGFP-SUN1^FL^ or EGFP-SUN2^FL^ with synthetic tetramethylrhodamine (TRITC)-labeled wild type (WT) KASH peptides from the mouse KASH protein nesprin-2 (TRITC-KASH2^WT^). Taken together, these results strongly suggest that our CFE-synthesized FL SUN proteins were directionally reconstituted in the ANMs such that their C-terminal KASH peptide-binding LDs were solvent-exposed and presumably homo-oligomerization competent.

Given that the orientation of SUN proteins reconstituted in ANMs was not random, we hypothesized that the CFE reaction lysates contain ER-derived microsomes that are capable of the co-translational translocation of membrane proteins^17^. Consistent with this hypothesis, we observed that the lipophilic membrane dye 1,1’-dioctadecyl-3,3,3’,3’-tetramethylindocarbocyanine perchlorate (DiI) stained small vesicle-like puncta present within our CFE reactions^8^. These puncta exhibited EGFP fluorescence that was presumably produced by the CFE synthesized EGFP-tagged FL SUN proteins present in these reactions. While we were unable to observe a clear co-localization of EGFP and DiI on these puncta due to their fast diffusion and the temporal limitations of our imaging system, we did observe co-localization between DiI and EGFP in puncta that had settled down onto the coverslip. These results suggested to us that our FL EGFP-tagged SUN protein constructs are inserted into the ER-derived microsomes present within the HeLa cell extracts used for their synthesis by CFE. They are also consistent with the observation of cellular membrane fragments present in CFE lysates obtained through the mechanical lysis of mammalian cells^18^. In fact, past studies identified the presence of ER fragments called microsomes in the lysates of Chinese hamster ovary cells used for CFE, which are translationally active and capable of the co-translational insertion of ER-targeted proteins in the microsomal membranes^19–21^.

The use of ER-derived microsomes in *in vitro* translation systems is a standard biochemical technique that enabled the discovery of translocons and membrane protein chaperones^22–24^. However, the proteins translated in such reactions were primarily secreted proteins (e.g. prolactin) that contain known ER-targeting signal sequences at their N-termini^25–27^. SUN proteins are known to have multiple regulatory elements at their N-termini, which result in them being targeted to the INM^28,29^ Thus, it is possible that a mammalian CFE system made from HeLa cells could have ER fragments that can recognize and translocate SUN proteins. Nevertheless, the mechanism underlying the reconstitution of CFE-synthesized FL membrane proteins in SUPER templates requires further investigation.

Our ability to successfully reconstitute CFE-synthesized FL SUN proteins in SUPER templates motivated us to explore the generality of expressing other membrane proteins and the possibility of co-reconstituting multiple membrane proteins in supported lipid bilayers. Having established ANMs as a facile HeLa-based CFE system that enables the efficient reconstitution of FL, KASH-binding competent SUN proteins in supported lipid bilayers *in vitro*, we sought to use them to experimentally probe three proposed regulators of LINC complex assembly: calcium (Ca^2+^) ions; redox environment; and the coiled-coil (CC)-containing domains present within the SUN protein LDs. Below, we will briefly discuss what is known about how these three potential regulators influence the assembly of functional LINC complexes.

Ca^2+^ ions are thought to influence LINC complex assembly in two ways. First, structural studies revealed the presence of a well-defined cation loop in the SUN domain of SUN2 that surrounds and coordinates a bound cation via five backbone carbonyls^13^. Importantly, the purified LD of SUN2 lost its ability to interact with KASH peptides when the cation loop was shortened by the deletion of one amino acid (i.e. ΔN600 in human SUN2). This result shows that the cation loop, and by extension the cation coordinated by the loop, are important for LINC complex assembly. Based on its coordination sphere, bond distances, and temperature factor, the bound cation in the cation loop of the previously reported crystal structure of human SUN2 was predicted to be a potassium ion. However, the potential malleability of the loop, the evolutionary relationship between SUN2 and the Ca^2+^ ion-binding F-lectin proteins, and the ionic environment of the NE lumen strongly suggest that the SUN protein cation loop may coordinate a Ca^2+^ ion in cells. Second, molecular dynamics modeling predicts that a Ca^2+^ ion interacts with residue E452 in the first of two CC-containing domains (i.e. CC1) present within the LD of human SUN2^30^. This residue was proposed to be involved in the monomer-trimer transition of the SUN2^LD^ by mediating the association of the SUN domain with the CC-containing regions of the SUN2^LD^ through Ca^2+^ ion-binding. Thus, Ca^2+^ ions are hypothesized to inhibit the homo-trimerization of SUN2 and presumably its ability to interact with KASH peptides. To date, this hypothesis has not been tested experimentally.

Regarding the potential effect of the redox environment of the NE lumen on LINC complex assembly, two disulfide bonds (DSBs) were detected in the previously reported crystal structure of a truncated LD construct of human SUN2 (amino acids 522-717) in complex with the KASH peptides of nesprin-1 or nesprin-2^13^. An intramolecular DSB that defines the arrangement of the cation loop of SUN2 is formed between two conserved cysteines (C601 and C705) and an intermolecular DSB is formed between a conserved cysteine (C563) of human SUN2 and a conserved cysteine residue (C-23) present within the KASH peptides of human nesprin-1/2. While the intermolecular SUN-KASH DSB appeared to be dispensable for the physical interaction of SUN2 and KASH1/2^13^, it was shown to be important for maximal mechanotransmission by SUN2 *in silico*^31^. This DSB is also important for actin-dependent rearward nuclear during centrosome orientation in fibroblasts polarizing for directional migration as well as nuclear anchorage in the developing hypodermis of *Caenorhabditis elegans*^31,32^. While less is known about the role of DSBs during the assembly of SUN1-containing LINC complexes, the conserved intermolecular SUN-KASH disulfide bond-forming cysteine residue found in SUN2 is also present in SUN1^13^. In addition, the SUN1 tertiary structure appears to involve interchain SUN1^LD^-SUN1^LD^ disulfide bonds that may promote the formation of higher-order homo-oligomeric complexes^33^, such as those predicted *in silico*^34^ and observed by fluorescence fluctuation spectroscopy within the NE of living cells^35,36^. Based on these results, DSBs may be potential targets for the regulation of LINC complex assembly and/or function. However, mechanistic experimental support for this hypothesis is currently lacking.

Lastly, we will briefly discuss how the CC-containing domains present within the LD of SUN proteins may regulate LINC complex assembly by controlling their homo- and/or hetero-oligomerization. *In vitro* data suggests that the CC-containing domains found within the LDs of SUN proteins may act as intrinsic dynamic regulators of their homo-oligomeric state and therefore ability to interact with KASH proteins^37^. The two CC-containing regions of the mouse SUN2^LD^ were shown to adopt distinct oligomeric states *in vitro* in solution, with CC1 and CC2 being monomeric and trimeric, respectively. Interestingly, the structure of a fragment of the mouse SUN2^LD^ that spans from CC2 to the SUN domain revealed that CC2 folds as a three-helix bundle referred to as the autoinhibitory domain that interacts with the SUN domain and traps it in an inactive conformation that cannot interact with KASH proteins. In contrast, CC1 was found to be a trimeric CC for the trimerization and activation of the SUN2 SUN domain. Recently, the mouse SUN1^LD^ was shown to also contain an autoinhibitory domain preceding its SUN domain that interacts with and inactivates the SUN domain^38^. Lastly, previous cell-based studies demonstrate that SUN1 and SUN2 can undergo hetero-oligomerization via a process that is mediated by both the SUN domain and the adjacent CC-containing regions^33,39–41^. At present, the physiological relevance of these CC-mediated regulations of SUN proteins remains unclear, as demonstrated by the fact that the CC domain present in the LD of the *C. elegans* SUN protein SUN-1 appears to be dispensable for functional LINC complex formation and successful meiotic chromosome pairing^42^.

In this work, we will further explore the mechanism underlying the reconstitution of CFE-synthesized FL membrane proteins in SUPER templates and present hypothesis for the observed membrane association of such proteins. This information is crucial for our ability to properly interpret the results of experiments performed using ANMs. Our work will also leverage our initial successes with the ANM platform to mediate the reconstitution of additional mammalian membrane proteins such as voltage-gated ion channels using the same approach with cell-free synthesis and appropriate choice of membrane lipids. The experimental approach described here provides a simple and powerful experimental platform for performing structure/function analyses of any membrane protein, nuclear or otherwise.

## RESULTS

### FL SUN proteins localize to ER-derived microsomes in HeLa cell-based CFE reactions

We previously investigated the reconstitution of CFE-synthesized FL SUN proteins in SUPER templates^8^. We observed that our EGFP- and 6x histidine (His_6_)-tagged FL SUN1 (EGFP-SUN1^FL^-His_6_; referred to as EGFP-SUN1^FL^ from here out) and SUN2 (EGFP-SUN2^FL^-His_6_; referred to as EGFP-SUN1^FL^ from here out) protein constructs were oriented in ANMs with solvent-exposed C-terminal KASH-binding SUN domains. We then demonstrated that both constructs retained the ability to interact with TRITC-KASH2^WT^ peptide *in vitro.* These results led us to explore the mechanism underlying FL SUN protein reconstitution in ANMs. We did this by using a standard Airfuge-based ultracentrifugation workflow (**Fig. 1B**), where we spun down CFE reactions with saturated yields (i.e. reactions that reached completion) of EGFP-SUN1^FL^, EGFP-SUN2^FL^, or soluble EGFP and tested their presence in equal volumes of supernatant and pellet fractions by in-gel fluorescence imaging (**Fig. 1C**; Full gel images are shown in **Fig. S1**). Since the volume of our CFE reactions was small (~10 μl), no significant membrane pellet was visible after ultracentrifugation. However, EGFP fluorescence was clearly visible in the pellet fractions, whereas no signal was detected in the supernatant fractions for any of the samples except for the soluble EGFP construct. Furthermore, the expression of the bacterial channel protein mechanosensitive channel of large conductance (MscL) that is capable of self-insertion into lipid membranes^43–45^ was also enriched in the pellet relative to the supernatant (**Fig. 1C**).

Staining of the CFE reactions with the ER tracker dye boron-dipyrromethene (BODIPY™)-FL glibenclamide revealed the existence of dense punctate structures in the lysate that were absent in the supernatant following ultracentrifugation (**Fig. 1D**). The presence of microsomal structures in our CFE lysates is further supported by cryo-electron microscopy (cryo-EM) (**Fig. 1E**). Lipid bilayer vesicles in the size range of 50-200 nm were clearly visible in our homemade CFE lysates, although their surface was not as densely covered with ribosomes as has been shown for canine pancreatic ER-derived microsomes *in situ* by cryo-EM tomography^46^. The results presented above are consistent with the translocation of FL SUN proteins in ER-derived microsomes.

### The direct reconstitution of CFE-synthesized FL mammalian membrane proteins in SUPER templates is mediated by endogenous ER-derived microsomal structures present in CFE reactions

Given the presence of endogenous ER-derived microsomes in our HeLa cell-based CFE reactions, we sought to see if we could enrich these reconstituted FL membrane proteins in ER-derived microsomes by ultra-centrifugation. Thus, we developed our Airfuge-based fractionation assay further by washing the pellet fraction after each centrifugation step to remove any proteins that were not associated with microsomes. In this assay, we recovered half of the supernatant after each step and diluted the pellet fraction in a 1:1 volume ratio before centrifuging the mixture again. After enriching for the microsomal fraction, we investigated whether we could directly reconstitute FL membrane proteins by introducing the microsomal fraction to SUPER templates designed to partially mimic the lipid composition of the budding yeast NE with added cholesterol^8,47^.

SUPER templates are made by mixing small unilamellar vesicles (SUVs) containing negatively charged lipids under conditions of high ionic strength with 5 μm diameter silica beads, which results in the fusion of the SUVs around the bead (**Fig. 2A**). This leads to the formation of supported lipid bilayers with larger areas than the surface of the silica bead such that a thin aqueous layer exists between the bead surface and the lipid bilayer^48^. This aqueous layer provides sufficient spacing between the bead and the lipid bilayer to accommodate membrane proteins with large soluble domains. In addition, fluorescence-based quantification previously demonstrated that the presence of excess membrane on SUPER templates minimizes the effects of substrate interaction^48^. Thus, SUPER templates serve as a superior model membrane system as compared to tightly substrate-bound supported lipid bilayers for FL SUN protein reconstitution.

**Figure 2.**
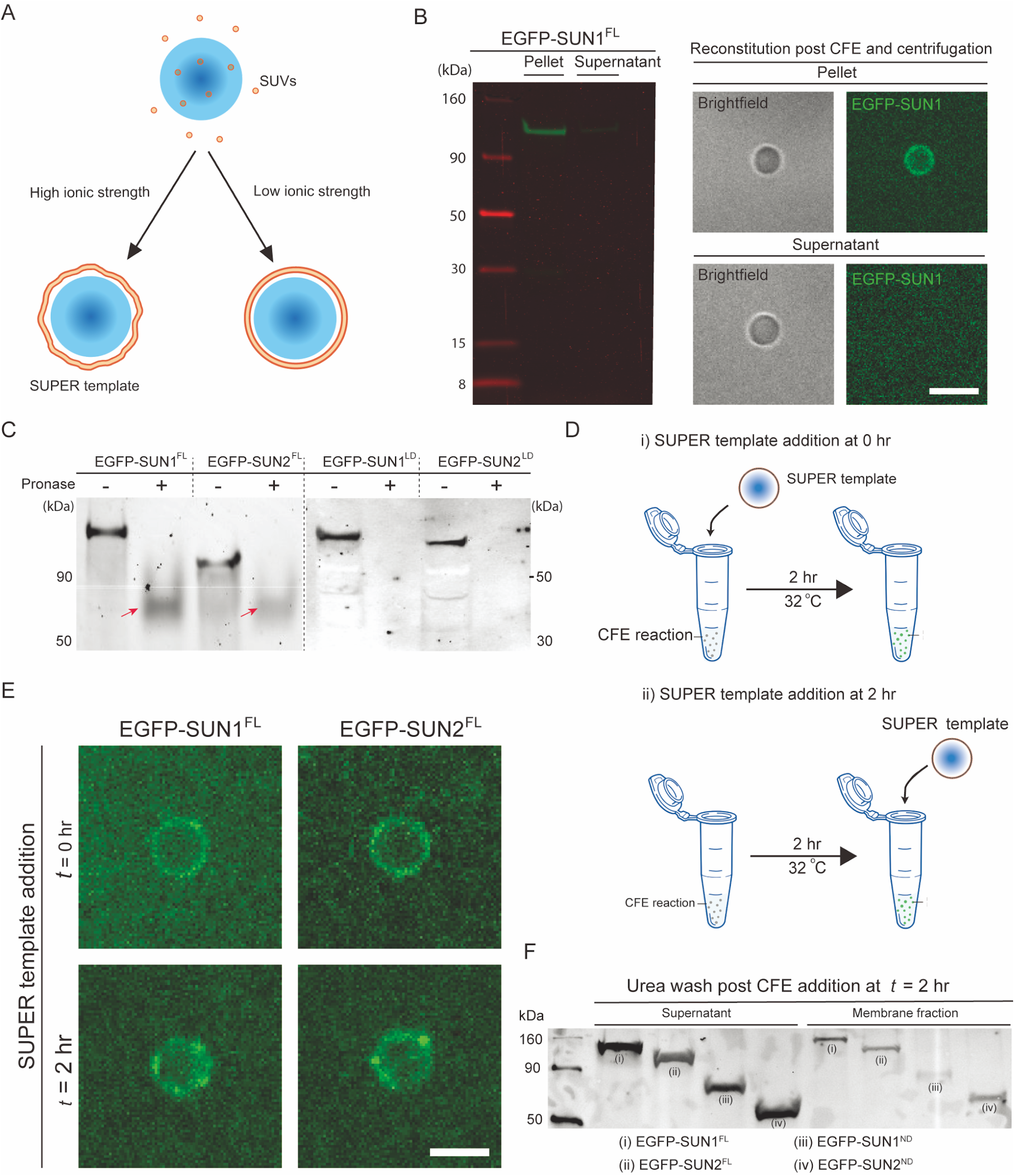
CFE synthesized FL SUN proteins are incorporated into SUPER templates via a potential fusionbased mechanism. A) Schematic depicting the formation of supported lipid bilayers on silica beads in the presence or absence of low or high ionic strength buffers. As shown, SUPER templates are typically created when the fusion of SUVs results in the formation of loosely bound unilamellar membranes under high salt conditions. B) In-gel fluorescent image of pellet and supernatant fractions of a CFE reaction with SUN1-EGFP. The pellet and supernatant fractions were isolated using the Airfuge fractionation assay and subsequently added to SUPER templates for imaging (right) or directly run on gel (left). Scale bars: 5 μm. C) In-gel fluorescence image of CFE reactions expressing the indicated EGFP-tagged proteins that were or were not exposed to pronase. Arrows: Incompletely pronase-digested proteins. D) Schematic illustrating how the ability of EGFP-tagged full-length SUN proteins to be reconstituted in SUPER templates added at the start of the CFE reaction (i) or 2 hours after its initiation (ii). The SUPER templates were washed in 1X PBS with 6M urea before imaging. E) Representative confocal fluorescent images of SUPER templates added to the indicated EGFP-tagged SUN protein CFE reactions at the indicated time points. Scale bars: 5 μm. F) Western blot of the supernatant and membrane fractions of CFE reactions to which SUPER templates were added 2 hours after initiating the synthesis of the indicated EGFP-tagged SUN protein constructs, as described in panel D(ii). The supernatant and membrane fractions represent SUN proteins in bulk CFE reactions and SUPER templates, respectively.

To ascertain the enrichment of these membrane proteins in the CFE pellet fractions, we performed SDS-PAGE analysis and confirmed that the pellet fractions had significantly more EGFP-SUN1^FL^ than the supernatant CFE fractions did after being washed (left panel, **Fig. 2B**). Consistent with our hypothesis that the direct reconstitution of CFE-synthesized mammalian membrane proteins on SUPER templates is mediated by endogenous ER-derived microsomal structures present in our CFE reactions, we observed EGFP fluorescence from the membranes of the SUPER templates that were incubated with the pellet fraction and not with the supernatant fraction of our centrifuged EGFP-SUN1^FL^ CFE reaction (right panel, **Fig. 2B**), suggesting that membrane protein incorporation into the SUPER templates is likely mediated by the fusion of ER-derived microsomes with the supported lipid bilayers.

To control for non-specific interactions between the CFE-synthesized proteins and the SUPER templates, we asked if the soluble green fluorescent genetically encoded Ca^2+^ indicator for optical imaging (G-GECO)^49^ behaved similarly to the SUN1 and SUN2 constructs tested above. Indeed, the pellet fraction contained some amount of G-GECO from the residual supernatant buffer CFE fraction (**Fig. S2**), indicating that this assay simply dilutes non-microsome-associated proteins.

### Fusion between ER-derived microsomes present in CFE reactions and SUPER templates enables the directional reconstitution of CFE-synthesized FL SUN proteins in these model membranes

To confirm the membrane insertion of EGFP-SUN1^FL^ and EGFP-SUN2^FL^ in ER-derived microsomes following their synthesis in CFE reactions, we performed a protease protection assay followed by an in-gel fluorescence measurement. The correct folding and insertion of FL SUN proteins requires them to span a lipid bilayer with their N- and C-termini on opposite sides of the membrane. Given that our FL SUN1 and SUN2 constructs are N-terminally tagged with EGFP, we were able to probe for EGFP fluorescence post digestion with the protease pronase. If the N-terminal nucleoplasmic domain (ND) of EGFP-SUN1^FL^ or EGFP-SUN2^FL^ reconstituted in SUPER templates were solvent exposed upon CFE synthesis, one would expect the EGFP to be degraded by the addition of pronase. As controls for pronase activity, we also generated in-gel fluorescence images of N-terminally EGFP-tagged constructs that encode the C-terminal LD of either SUN1 (EGFP-SUN1^LD^) or SUN2 (EGFP-SUN2^LD^) following their synthesis by CFE and exposure to protease (**Fig. 2C**; full gel images are shown in **Fig. S3A**).

In the absence of pronase (gel lanes labeled “-“;**Fig. 2C**), we observed fluorescent bands at the sizes predicted for our constructs (e.g. EGFP-SUN1^FL^: ~130 kD; EGFP-SUN2^FL^: 110 kD; EGFP-SUN1^LD^: ~80 kD; and EGFP-SUN2^LD^: ~80 kD). In contrast, after being digested with pronase (gel lanes labeled “+”; **Fig. 2C**), we no longer observed fluorescent bands at the same size as in our non-digested samples. Specifically, we detected the presence of much smaller bands (red arrows; **Fig. 2C**) for EGFP-SUN1^FL^ or EGFP-SUN2^FL^ and the complete absence of fluorescent bands for EGFP-SUN1^LD^ or EGFP-SUN2^LD^ (**Fig. 2C**). Based on these results, we can conclude that a fraction of our EGFP-tagged FL SUN proteins is successfully translocated into the ER-derived microsomes present in our CFE reactions with their N-terminal NDs being solvent exposed.

We then asked how the topology of SUN proteins reconstituted in SUPER templates occurs in a unidirectional manner, given their translocation into ER-derived microsomes during their synthesis by CFE. One plausible hypothesis is that SUN proteins are directionally reconstituted in SUPER templates by the spontaneous fusion of ER-derived microsomal membranes with the supported lipid bilayers of the SUPER templates. Previous studies showed that cations and the electrostatic charges of lipids are key determinants of spontaneous vesicle fusion, especially between liposomes with high curvature and planar lipid bilayers^50,51^. Since CFE reactions contain mM levels of potassium and magnesium ions and SUPER templates are made with charged lipids (i.e. phosphatidylserine) and lipids that mediate spontaneous fusion at room temperature (i.e. phosphatidylethanolamine)^52,53^, the mechanics of membrane fusion can ensure a consistent topology of reconstituted integral membrane proteins as opposed to direct membrane insertion.

To probe this potential mechanism of CFE-synthesized FL SUN protein reconstitution in SUPER templates, we compared the SUPER template localization of EGFP-SUN1^FL^ and EGFP-SUN2^FL^ under two conditions. In the first (**Fig. 2Di**), the SUPER template is added to the CFE reactions at the beginning of the reaction. In the second (**Fig 2Dii**), the SUPER template is added to the CFE reactions at the end of the reaction. If the EGFP-tagged FL SUN protein constructs were co-translationally inserted into the SUPER templates, a significant reduction in the levels of SUPER template-associated EGFP fluorescence would be expected for the second condition relative to the first. However, we observed a similar association of both EGFP-SUN1^FL^ and EGFP-SUN2^FL^ with the SUPER templates under both conditions (**Fig. 2E**). This result indicates that the reconstitution of CFE-synthesized FL SUN proteins in SUPER templates can be carried out by the addition of SUPER templates to saturated CFE reactions. While the direct post-translational insertion of FL SUN proteins in the SUPER templates may explain the lack of an observable difference between the two conditions described above, it is unlikely that SUN proteins with large soluble domains and multi-membrane-associating domains can self-insert into synthetic lipid bilayers while maintaining their specific membrane topologies. Rather, vesicle fusion is the most plausible mechanism of CFE-synthesized FL membrane protein reconstitution in SUPER templates.

To further explore the validity of vesicle fusion as the main mechanism underlying membrane protein reconstitution in SUPER templates, we tested the ability of the chaotropic agent urea to disrupt the SUPER template-association of EGFP-SUN1^FL^ and EGFP-SUN2^FL^. We also tested the impact of urea on the SUPER template-association of two EGFP-tagged SUN protein ND constructs EGFP-SUN1^ND^ and EGFP-SUN2^ND^, which we previously demonstrated were peripherally associated with the outer leaflets of SUPER templates^8^. Consistent with a urea wash being able to disrupt weak associations of proteins with membranes^54,55^, we observed that all four of the constructs we tested exhibited strong interactions with the SUPER templates as demonstrated by their presence in the membrane fractions as well as the supernatant fractions (**Fig. 2F**; full gel images are shown in **Fig. S3**). The supernatant fractions described here correspond to the bulk CFE reactions that were added to the SUPER templates at the maximal levels (i.e. completed reactions) of the indicated CFE-synthesized EGFP-tagged FL SUN protein constructs. Altogether, the results presented here suggest that our HeLa cell-based CFE system enables the translocation of FL membrane proteins into ER-derived microsomes that are subsequently directionally reconstituted in SUPER templates most likely through vesicle fusion.

### Ca^2+^ ions and redox state are critical regulators of SUN-KASH interactions *in vitro*

Armed with a better understanding of how CFE-synthesized FL SUN proteins are directionally reconstituted in SUPER templates, we next tested the impact of the chemical environment (e.g. Ca^2+^ ions and redox state) on the ability of FL SUN proteins to interact with KASH peptides. To test the potential role for Ca^2+^ ions in promoting LINC complex assembly, we took advantage of the negligible levels of Ca^2+^ ions present in our CFE reactions^56,57^. More specifically, we asked if the addition of 1 mM calcium chloride (CaCl_2_) positively or negatively impacted the ability of SUPER templates containing reconstituted CFE-synthesized EGFP-SUN1^FL^ or EGFP-SUN2^FL^ to recruit TRITC–KASH2^WT^. Similarly, we asked if either of these CFE-synthesized FL SUN protein-containing SUPER templates were impaired or enhanced in their ability to recruit synthetic TRITC-labeled nesprin-2-derived KASH peptides lacking the four C-terminal amino acids required for the SUN-KASH interaction to normally occur (TRITC-KASH2^ΔPPPT^)^58^. We observed that the addition of CaCl_2_ significantly increased the ability of ANMs containing reconstituted CFE-synthesized EGFP-SUN1^FL^ or EGFP-SUN2^FL^ to recruit TRITC-KASH2^WT^ above background levels measured in the absence of CaCl_2_ (**Figs. 3A-D**). In contrast, the recruitment of TRITC-KASH2^ΔPPPT^ to ANMs harboring either CFE-synthesized EGFP-SUN1^FL^ or EGFP-SUN2^FL^ was significantly reduced relative to the recruitment of TRITC-KASH2^WT^ to either type of AMN (**Figs. 3A-D**). Taken together, these results support the hypothesis that Ca^2+^ ions promote SUN-KASH interactions.

**Figure 3:**
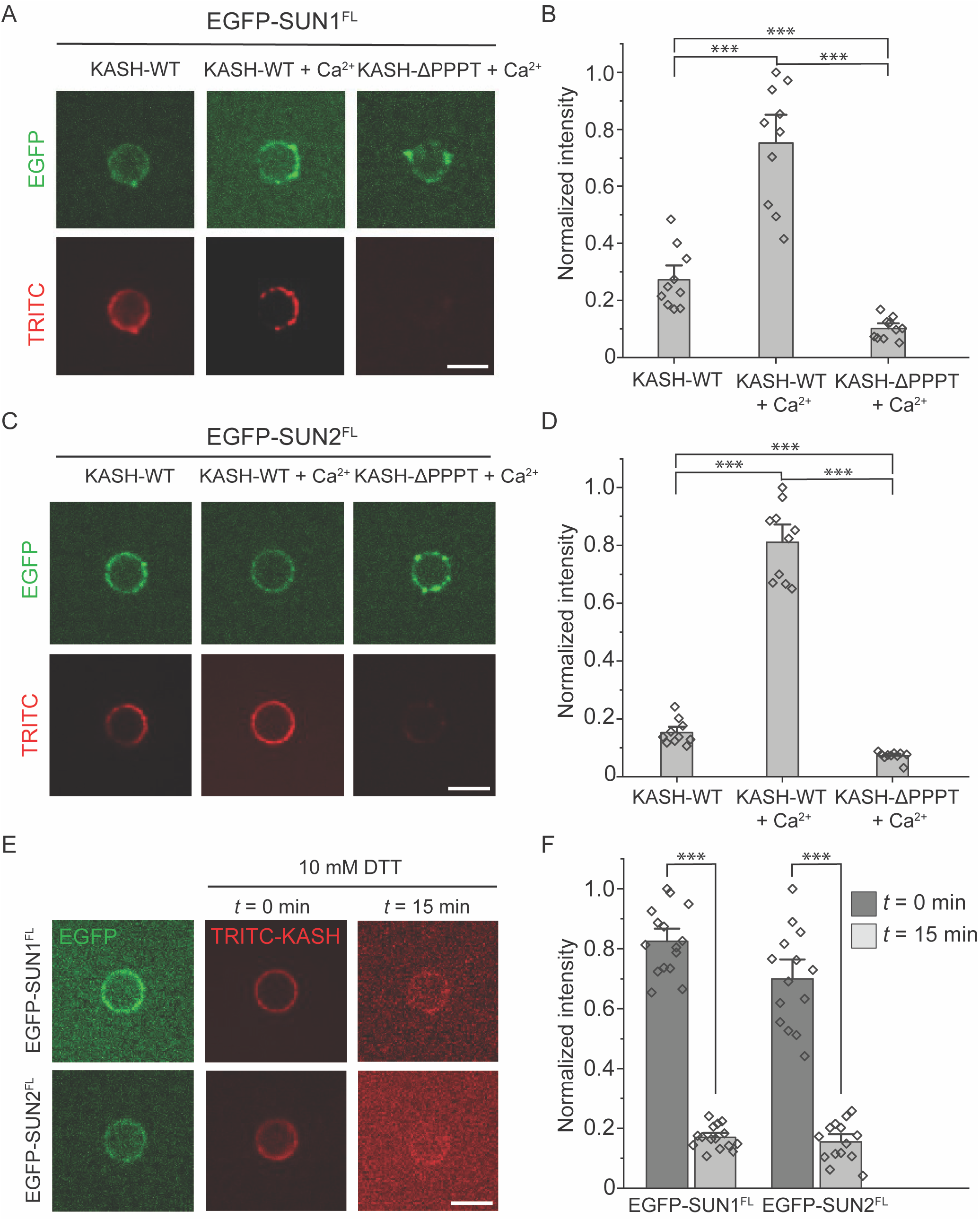
The SUN1/2-KASH2 interaction is enhanced by Ca^2+^ ions and stabilized by oxidizing conditions. A and C) Representative confocal fluorescence images of SUPER templates containing the indicated reconstituted EGFP-tagged FL SUN protein constructs after incubation with the indicated TRITC-labeled KASH2 peptides in the presence and absence of 1 mM CaCl_2_. B and D) Bar graphs depicting the normalized intensity of the indicated TRITC-labeled KASH2 peptides recruited to the SUPER templates containing the indicated EGFP-tagged FL SUN protein constructs in the absence or presence of 1 mM CaCl_2_. E) Representative confocal fluorescence images of SUPER templates containing the indicated reconstituted EGFP-tagged FL SUN protein constructs following incubation with 10 mM DTT. F) Bar graph depicting the normalized intensity of the indicated TRITC-labeled KASH2 peptides recruited to the SUPER templates containing the indicated EGFP-tagged FL SUN protein constructs following incubation with 10 mM DTT. Scale bars: 5 μm. Error bars represent standard error of the mean. 10-15 beads were analyzed for each condition from n = 3 experiments. ***: *p* < 0.001 by pairwise Student’s *t*-test.

We next used this experimental platform to investigate the effect of redox state on LINC complex assembly. Specifically, we tested the effect of high concentrations (e.g. 10 mM) of the strong reducing agent dithiothreitol (DTT) on pre-formed SUN-KASH interactions using our *in vitro* experimental platform. We found that exposing either CFE-synthesized EGFP-SUN1^FL^- or EGFP-SUN2^FL^-containing SUPER templates bound to TRITC-KASH2^WT^ for 15 minutes significantly reduced the amount of bound TRITC-KASH2^WT^ relative to the amount present at the start of the experiment by 4-fold (**Figs. 3E-F**). In contrast, no significant decrease in SUPER template-associated EGFP fluorescence was detected at the end of these experiments (data not shown). Taken together, these results identify Ca^2+^ ions and redox state as key regulators of SUN-KASH interactions *in vitro*. They also demonstrate the ease with which our HeLa cell-based CFE and SUPER template system can be used to investigate mechanistic hypotheses regarding LINC complex assembly and regulation that may be difficult to address in cells.

### SUN protein homo- and hetero-oligomerization is mediated by their luminal CC domains

To further test the hypothesis that the CC domains of SUN proteins act as intrinsic dynamic regulators of SUN protein homo- and hetero-oligomerization, we asked if reconstituted CFE-synthesized FL SUN proteins directionally reconstituted in SUPER templates could form homo- or hetero-oligomers with soluble fragments of their LDs. To do this, we used CFE to synthesize SUN1^FL^ and SUN2^FL^ in the presence of the FluoroTect™ Green Lys *in vitro* translation labeling system (referred to as GreenLys from here on). GreenLys enables the fluorescent labeling and detection of *in vitro* synthesized proteins via a lysine-charged tRNA labeled with the fluorophore BODIPY™-FL at its ε position. The reconstituted CFE-synthesized GreenLys-labeled FL SUN proteins in SUPER templates (**Fig. 4A**). Next, we designed an assay to investigate the interaction of reconstituted WT SUN proteins with their soluble counterparts (i.e. their CFE-synthesized LDs) (**Fig. 4B**), where the same SUPER templates were exposed to two rounds of CFE reactions expressing different proteins. The SUPER templates were washed in between each step.

**Figure 4:**
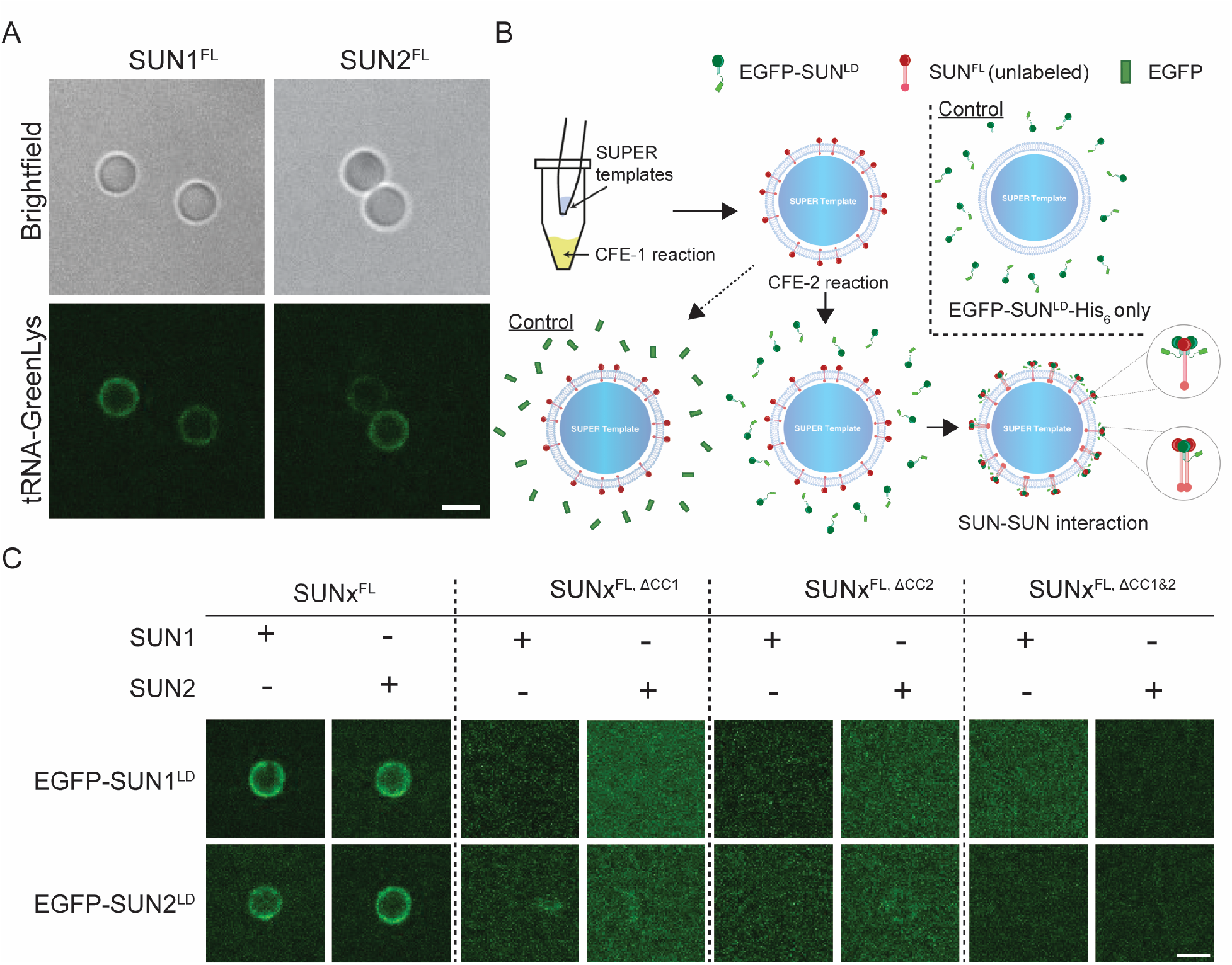
The luminal CC-containing domains are important for homo- and hetero-typic SUN protein interactions. A) Representative confocal fluorescence images of SUPER templates containing the indicated reconstituted CFE synthesized FL SUN protein constructs. B) Schematic depicting the workflow for testing the ability of CFE synthesized EGFP-tagged SUN protein LD constructs to be recruited to SUPER templates containing reconstituted CFE synthesized FL SUN protein constructs. To control for non-specific protein-protein interactions, the recruitment of soluble CFE synthesized EGFP to SUPER templates containing reconstituted CFE synthesized FL SUN protein constructs was be tested (bottom left). To control for non-specific interactions between the SUPER template and the CFE synthesized EGFP-tagged SUN protein constructs, the ability of these constructs to be recruited to SUPER templates lacking any reconstituted CFE synthesized FL GreenLys-labeled SUN protein constructs will be tested (top right). C) Representative confocal fluorescence images of SUPER templates containing the indicated reconstituted CFE synthesized FL SUN protein constructs and their ability to recruit the indicated CFE synthesized EGFP-tagged LD SUN protein constructs. Experiments were repeated 3 times. Scale bars: 5 μm.

In the first round of CFE and SUPER template reconstitution, CFE-synthesized GreenLys-labeled SUN1^FL^ and SUN2^FL^ were reconstituted in SUPER templates, which were then treated with a second round of CFE synthesizing either EGFP or EGFP-SUN1/2^LD^. Expressing EGFP-SUN^LD^ without expressing SUN^FL^ in the first CFE reaction served as a control (Fig. 4B). As we previously reported^8^, neither EGFP-SUN1^LD^ nor EGFP-SUN2^LD^ strongly associated with the SUPER templates in the absence of directionally reconstituted CFE-synthesized GreenLys-labeled SUN1^FL^ or SUN2^FL^. Therefore, any EGFP fluorescence detected on the SUPER templates after the second round of CFE with EGFP-SUN1^LD^ or EGFP-SUN2^LD^ indicates the presence of an interaction between the reconstituted CFE-synthesized GreenLys-labeled FL SUN protein and its corresponding soluble EGFP-tagged LD.

Additionally, given two independent rounds of CFE, it is possible to expose GreenLys-labeled SUN1^FL^ to EGFP-SUN2^LD^ and GreenLys-labeled SUN2^FL^ to EGFP-SUN1^LD^, in order to study heteromeric SUN protein interactions. Thus, we created CC domain deletion mutants for both CFE-synthesized GreenLys-labeled FL SUN proteins in which either or both of the CC domains were deleted (i.e. SUN1/2^FL,ΔCC1^, SUN1/2^FL,ΔCC2^, SUN1/2^FL,ΔCC1&2^). The membrane association and SUPER template reconstitution of each of the CFE-synthesized GreenLys-labeled FL SUN protein CC deletion mutant constructs was tested using GreenLys-labeling as described above (**Fig. S4**).

We carried out all combinations of the two CFE reactions with FL WT SUN proteins or FL SUN protein mutants that lacked CC1, CC2, or both CC1 and CC2 (**Fig. 4C**). We found that the EGFP fluorescence detected after the second round of CFE of the EGFP-tagged SUN1^LD^ or SUN2^LD^ constructs was only observed on the SUPER templates when the CFE-synthesized GreenLys-labeled WT SUN1^FL^ or SUN2^FL^ constructs were reconstituted in the first CFE reaction. CFE of soluble EGFP alone in the second round served as a control for the non-specific interaction of EGFP with the SUPER templates (**Fig. S5**). Hence, for reconstituted CFE-synthesized GreenLys-labeled WT SUN1^FL^ or SUN2^FL^, we detected robust homo- and hetero-typic interactions between their LDs. In the case of the reconstituted CFE-synthesized GreenLys-labeled CC deletion SUN1 or SUN2 constructs, no significant fluorescence was observed on the SUPER templates for any of the experiments, indicating a lack of homo- or hetero-typic interactions between these SUN protein mutants and the EGFP-tagged LDs of SUN1 or SUN2. Collectively, our findings suggest that the deletion of either CC1 or CC2 from SUN1 or SUN2 is sufficient to disrupt both homo- and hetero-typic SUN protein interactions.

### Three different CFE-synthesized FL membrane proteins can be simultaneously reconstituted in a single SUPER template

Based on the results presented thus far, one might imagine that other mammalian membrane proteins with multiple membrane-spanning domains can be reconstituted using the same strategy undertaken for reconstitution of SUN proteins. Additionally, if the membrane protein reconstitution is mediated by microsome-fusion, multiple membrane proteins could be simultaneously reconstituted on a SUPER template by enriching the microsomal fraction of independent CFE reactions and depositing these fractions together on SUPER templates. To test this possibility, we carried out an experiment following the flowchart depicted in **Figure 5A** and used CFE to independently synthesize several additional membrane-spanning proteins, including the human codon-optimized channelrhodopsin 2 (hChR2) harboring a C-terminal monomeric Cherry and 6x histidine tag fusion (hChR2-mCherry-His_6_, referred to as hChR2-mCherry from here on), the inward rectifying potassium channel Kir2.1 harboring a C-terminal superfolder GFP fusion and 6x histidine tag fusion (Kir2.1-sfGFP-His_6_, referred to as Kir2.1-sfGFP from here on), and the voltage-gated potassium channel K_v_1.2 harboring a C-terminal 6x histidine tag (K_v_1.2-His_6_, referred to as K_v_1.2 from here on). Note that K_v_1.2 has a BFP tag, but proper detection requires a high expression level, so it is labeled with GreenLys unless otherwise noted.

**Figure 5.**
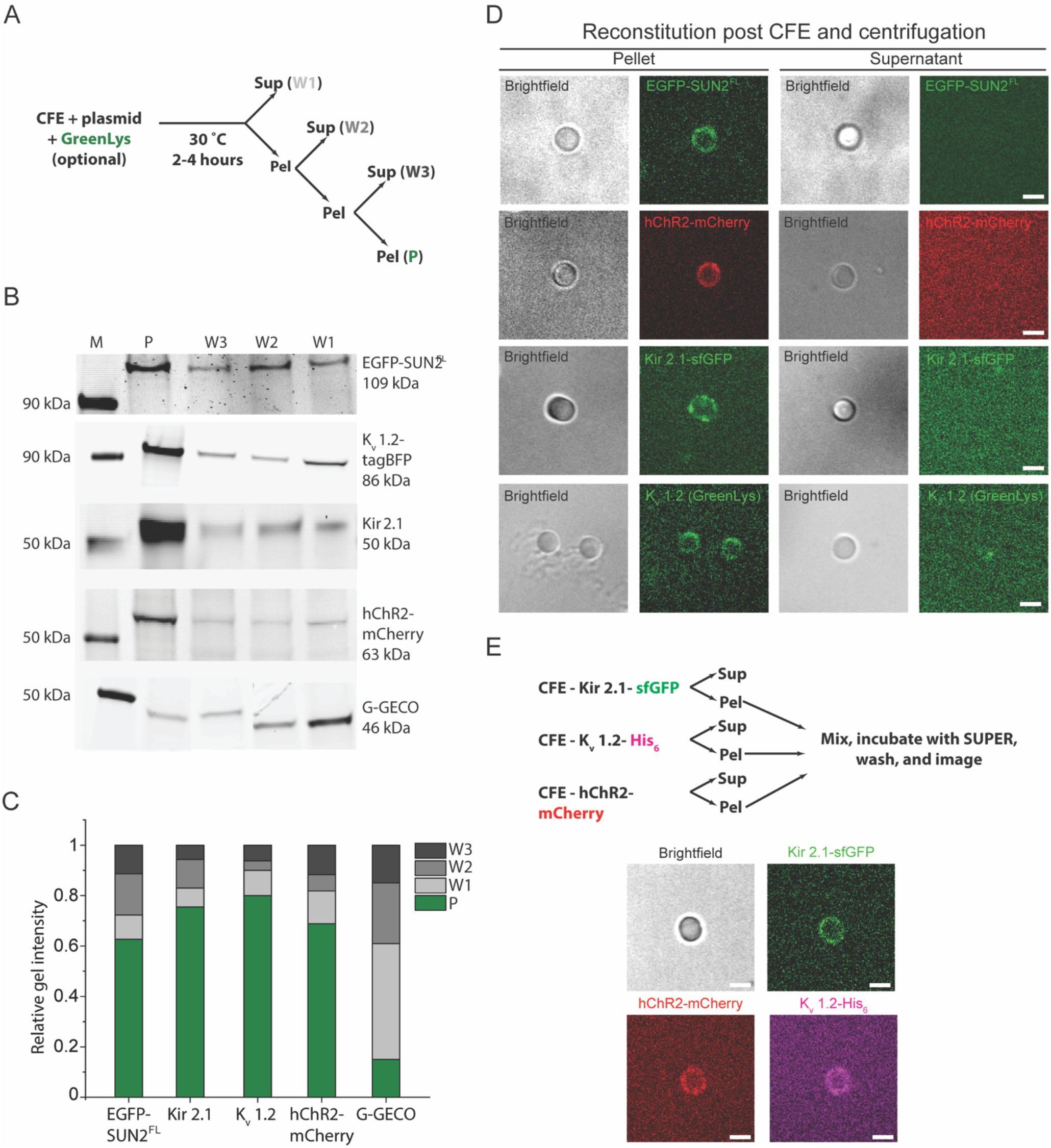
Direct reconstitution of FL membrane proteins mediated by microsome fusion is used for the reconstitution of different or multiple membrane proteins. A) Illustration of the experimental workflow used in panels B-E of this figure. B) Representative brightfield and confocal fluorescence images of SUPER templates incubated in and isolated from the pellet (denoted as P in panel A) or the supernatant fraction from the first wash (denoted as W1 in panel A) from microsome enrichment experiments where the indicated membrane protein constructs were synthesized. Scale bars: 5 μm. C) Representative in-gel fluorescence images of the pellet and supernatant fractions from the microsome enrichment experiment depicted in panel A for the indicated CFE synthesized and GreenLys-labeled protein constructs. Lane M indicates the protein ladder. D) Bar graph of the normalized band intensities quantified from the in-gel fluorescence images presented in panel D. E) Experimental design for multiple membrane protein reconstitution (top) and representative confocal fluorescence images of SUPER templates incubated with microsome-enriched fractions of completed CFE reactions that synthesized the indicated proteins (bottom). Scale bars: 5 μm.

The enrichment of these membrane proteins in the CFE pellet fractions, all synthesized with GreenLys incorporated, was determined by SDS-PAGE, which demonstrated that the pellet fractions had significantly more membrane proteins than the supernatant CFE fractions did after each wash while the supernatant CFE fractions lost progressively more synthesized membrane protein after each wash (**Figs. 5B-C**). Also shown are the bands from similar enrichment of EGFP-SUN2^FL^ after completion of bulk CFE in the absence of SUPER templates. Importantly, the soluble G-GECO control construct exhibited the opposite behavior from the four membrane proteins tested above. Next, we added the enriched pellet fractions for each CFE reaction using the above described Airfuge fractionation assay to SUPER templates that were made using ANM composition for EGFP-SUN2^FL^ and polar brain extract lipids for the ion channels, as this lipid composition resembles their native lipid environment^59,60^. Fluorescent confocal microscopy revealed that each membrane protein including EGFP-SUN2^FL^ was successfully reconstituted in the SUPER templates post CFE and centrifugation (**Fig. 5D**).

Since all the ion channels used in this experiment had a C-terminal 6x histidine tag, for reconstituting all three ion channels into the same lipid bilayer membrane, we did not introduce all the pellets at once to the SUPER template to avoid labeling the other CFE-synthesized ion channels. After incubating the fluorescent antihistidine tag antibody with the SUPER templates with K_v_1.2, they were washed and then incubated with the mixture of the Kir2.1-sfGFP- and hChR2-mCherry-containing CFE pellet fractions followed by additional washing steps. Using this approach, we were able to successfully reconstitute three different membrane proteins at once in a SUPER template (**Fig. 5E**). The demonstration further supports the possibility of a fusion-mediated mechanism for reconstitution of CFE-synthesized FL membrane proteins in SUPER templates.

## DISCUSSION

In this work, we demonstrate the use of a facile HeLa cell-based CFE system to study FL membrane protein biochemistry in supported lipid bilayers *in vitro.* We first investigated the mechanism underlying the directional reconstitution of CFE-synthesized FL membrane proteins in SUPER templates that we previously reported^8^. Next, we leveraged the specific topology of CFE-synthesized FL SUN proteins that were directionally reconstituted in SUPER templates to study hetero-typic SUN-KASH interactions as well as homo- and heterotypic SUN protein interactions.

The use of FL SUN1 and SUN2 labeled with EGFP or incorporated GreenLys and their reconstitution in SUPER templates containing 5 μm diameter silica beads enabled the direct detection of protein localization and subsequent protein-protein interactions by confocal fluorescence microscopy. With a bead-based platform such as the SUPER templates^48^, it was possible to design simple *in vitro* biochemical assays to probe protein function owing to the ease of handling the CFE-synthesized and reconstituted FL SUN proteins. The major advantage of a CFE system is ease and modularity in synthesizing many different proteins^61,62^. As a result, homo- and heterotypic SUN protein interactions could be probed through multiple rounds of CFE performed on the same batch of SUPER templates, while expressing different protein constructs. Another powerful advantage of this platform is its ability to simultaneously synthesize and translocate FL SUN proteins into the synthetic membranes of SUPER templates. While the C-terminally exposed topology of FL SUN1 and SUN2 in the SUPER templates was a discovery and not a controlled reconstitution, the evidence of ER-derived microsome-based reconstitution of SUN proteins in SUPER templates, potentially through membrane fusion, provides a way to possibly control the orientation of FL membrane reconstituted in SUPER templates, as previously reported^63,64^.

Unlike standard supported lipid bilayers, SUPER templates serve as an ideal membrane model in this study because they possess an excess membrane reservoir that is not tightly bound to the bead substrate^16^. This is reflected by the ability of reconstituted CFE-synthesized FL WT EGFP-tagged SUN1 or SUN2 to interact with synthetic TRITC-labeled KASH2^WT^ peptides, which should only be possible if the reconstituted SUN proteins present in the SUPER templates were homo-oligomerization competent. Further, the observation of small fluorescent puncta on the SUPER templates could be evidence of SUN protein clustering in these excess membrane reservoirs suggesting the presence of sufficient protein mobility, which is less readily achievable in conventional supported lipid bilayers^65^. The short timescale for the entire reconstitution step with sufficient yield of proteins to mediate fluorescence-based detection is a key feature of our HeLa cell-based CFE platform that enabled us to carry out the studies presented here.

Our results only provide a window into the range of experimental possibilities accessible with our HeLa cell-based CFE platform. For instance, one could look at the effect of different concentrations of Ca^2+^ ions on KASH binding to reconstituted CFE-synthesized FL SUN proteins. The specific cysteine residues responsible for DSB formation during KASH peptide-binding^32^ can be mutated and their effect studied without significant changes to the demonstrated approach. Further insights can be gained by the quantitative estimation of the timescale of DTT-based dissociation of the LINC complex. Similar extensions of the current work can be carried out to further investigate the possibility of hybrid LINC complexes based on hetero-typic SUN1-SUN2 interactions. Finally, the entire approach can be directly applied to the study of LINC complex formation between SUN1 or SUN2 with non-nesprin-2-derived mammalian genome-encoded KASH peptides, including nesprin-1, nesprin-3, nesprin-4, KASH5, and lymphoid restricted membrane protein (a.k.a. Jaw1)^15^. This is especially interesting in light of recent structural studies, which revealed that alternative binding modes exist for the KASH peptides derived from nesprin-3, nesprin-4, and KASH5 with SUN2 homo-trimers relative to the KASH peptides derived from nesprin-1 or nesprin-2^66^. Moreover, the same study demonstrated that a SUN2 homo-trimer could simultaneously interact with different KASH peptides. While it has been proposed that these differences may have an important role in regulating the SUN-KASH network, the underlying mechanisms remain poorly defined.

Overall, our findings provide strong support for some of the existing hypotheses about LINC complex assembly, while new insights were gained with respect to the importance of biochemical interactions in SUN-KASH binding. Specifically, we found that Ca^2+^ ions significantly enhance the ability of reconstituted FL SUN proteins to bind to KASH peptides, but are not necessary for LINC complex assembly. Since the ER is contiguous with the ONM, ER-mediated calcium signaling elicited by mechanical signals^67^ may strengthen hetero-typic SUN-KASH interactions, thus priming LINC complex-dependent mechanotransduction^68^. On the other hand, our data suggest that DSBs are indispensable for the maintenance of stable SUN1/2-KASH2 interactions. This result is consistent with the recent demonstration that the conserved interchain SUN-KASH DSB is required for actin-dependent nuclear anchorage and movement as well as LINC complex-dependent mechanotransmission^31,32^. By extension, the redox state of the ER lumen, which is closely linked to ER protein-folding homeostasis^69^, may be a key regulator of stable LINC complex assembly. Finally, it is likely that homo- and hetero-typic SUN protein interactions require the presence of CC domains within their LDs. Future efforts are needed to identify the exact DSBs that are important for the hetero-typic SUN-KASH interaction in this system. It will be informative to test the ability of FL SUN protein constructs that harbor mutations designed to inactivate either the intramolecular DSB known to be important for stabilizing the cation loop of SUN2 or the intermolecular SUN-KASH DSB.

In summary, we developed a modular platform for the study of *in vitro* reconstituted CFE-synthesized membrane proteins with an emphasis on the real-time detection through fluorescence microscopy and a possibility to use simple biochemical techniques to probe SUN protein biology. Such a platform can be extended to study the biochemical and biophysical properties of other difficult-to-study membrane proteins.

## METHODS

### Materials

#### Chemicals

All lipids were purchased from Avanti Polar Lipids. Other reagents used are previously described^8^. Dithiothreitol (DTT) was purchased from Sigma-Aldrich. FluoroTect™ Green Lys-tRNA was purchased from Promega.

#### Plasmids

All EGFP tagged and untagged full-length mouse SUN protein plasmids (UniProtKB – Q9D666 and Q8BJS4) were used in our previous study^8^. All CC mutants were generated from untagged WT SUN1 and SUN2 constructs by PCR-mediated mutagenesis (NEB site directed mutagenesis kit) (**Table S1**).

### Preparation of SUPER templates

SUPER template beads were made following a previously published protocol^64^ and our prior work with SUN proteins^8^. ANM composition consists of 45% DOPC, 27% DOPE, 9% DOPS, 2.2% phosphatidic acid, and 16.8% cholesterol. For reconstitution of ion channels, porcine brain polar lipid extract with the composition of 12.6 % PC, 33.1% PE, 4.1% PI, 18.5% PS, 0.8% PA, and 30.9% unknown lipids were used.

### CFE of SUN proteins

A homemade HeLa CFE system was made based on previously published protocols^54,55^. A typical reaction volume of 10 μl was assembled with 5 nM plasmid in a 1.5 mL microcentrifuge tube and incubated at 30°C for up to 3 hours. Most reactions reached saturation after 2 hours. For the standard reconstitution of SUN proteins (**Fig. 2Di**), SUPER templates were added to the CFE reaction prior to incubation. When SUPER templates were added to completed CFE reactions, the mixture was incubated at room temperature for approximately 15 minutes with intermittent shaking prior to subsequent analysis. All reconstituted SUPER templates were washed with 2 ml 1X PBS with or without 6 M urea, as indicated. For fluorescent visualization of non-EGFP-tagged proteins, GreenLys was added to the CFE reaction at a dilution of 1:50 as recommended in the manufacturer’s protocol.

### Ultracentrifugation with Airfuge

A Beckman Coulter Airfuge was used to spin down CFE reactions. Completed CFE reactions were collected in a 1.5-mL microcentrifuge tube and then mixed with 30 μl of extraction buffer (20 mM HEPES-KOH, pH 7.5, 45 mM potassium acetate, 45 mM KCl, 1.8 mM magnesium acetate, 1 mM DTT). Next, 40 μl of the mixture was transferred to an ultracentrifuge tube and centrifuged at around 100,000 x g for 15 minutes at room temperature using the Airfuge (Beckman Coulter). After centrifugation, 20 μl of the supernatant were carefully recovered so as not to disturb the pellet and transferred to a 1.5-mL microcentrifuge tube. The remaining 20 μl of pellet fraction was resuspended by pipetting the sample up and down before being transferring to another microcentrifuge tube. To investigate protein incorporation, 2 μl of SUPER templates were added and incubated with the supernatant or pellet fractions for 30 minutes and then centrifuged at 300 x g for 3 minutes. The SUPER templates were visible as a small white pellet, which was gently resuspended by tapping for downstream analysis.

### Protease digestion assay

Pronase (Roche Cat No.: 10165921001) was dissolved in de-ionized water at a stock concentration of 10 mg/ml. Next, 2 μl of this stock was added to 8 μl of a bulk CFE reaction and incubated at room temperature for 15-20 minutes. Thereafter, 2 μl of 25X Complete mini protease inhibitor cocktail (Millipore-Sigma Cat No.: 11836153001) was added to the digested CFE and incubated for 10 minutes. This mixture was subsequently analyzed by SDS-PAGE or Western blot.

### TRITC-KASH2 peptide-binding assay

Reconstituted SUN proteins on SUPER templates were washed with 1X PBS. The beads were resuspended in 30 μl of 1X PBS and distributed into 9 μl aliquots in Eppendorf tubes. To two aliquots, ~300 nM TRITC-KASH^WT^ was added followed by addition of 1 mM CaCl_2_ (final concentration) in one and an equal volume of DI water in another. To the third aliquot, ~300 nM of TRITC-KASH^ΔPPPT^ was added followed by dilution with CaCl_2_ to a final concentration of 1 mM. The three tubes were then incubated at room temperature for 10 minutes before subsequent imaging. In order to investigate the effect of DTT, 10 mM DTT (final concentration) was added to the sample after incubation of the TRITC-KASH2 peptide to the SUN protein-reconstituted SUPER templates.

### In-gel imaging and Western blots

In-gel imaging of SUN proteins was carried out on a Sapphire biomolecular imager (Azure Biosystems) for both EGFP-tagged and GreenLys-labeled proteins. Samples were loaded on 4-20% gradient bis-Tris PAGE gels (Bio-Rad) with 4X Laemmeli loading buffer (Bio-Rad). Samples were not heated to retain in-gel EGFP and GreenLys fluorescence. A rabbit polyclonal anti-GFP antibody at a dilution of 1:2000 (Abcam ab6556) was used to detect EGFP-tagged SUN mutants by Western blotting. For immunostaining his-tagged protein on SUPER template, a 1:2000 dilution of a mouse anti-Penta-His Alexa Fluor 647 conjugate (Qiagen) was used.

### Confocal fluorescence microscopy and analysis

Confocal images of SUPER templates were taken as previously described on a spinning disk confocal microscope and subsequent analysis was carried out using ImageJ as before^8^. Briefly, 5-10 line scans for each bead for the relevant scenario were taken and the peak intensities recorded. Subsequently, their mean was taken and combined with intensity values from 10-15 beads for background correction and statistical analysis. Pairwise student *t*-tests and one factor ANOVA for the plots obtained in Fig. 3 were carried out from background subtracted intensity values obtained using ImageJ. The values were then normalized by the maximum intensity of the group before plotting.

### Cryo-EM grid preparation

CFE reactions expressing EGFP-SUN1^FL^ were ultracentrifuged as mentioned above and 3.5 μL of the pellet fraction was applied to glow discharged grids (Quantifoil R1.2/1.3 Cu 200 mesh, glow discharged 1 minute, 5 mA) and frozen by plunging into liquid ethane using an FEI Vitrobot Mark IV (100% humidity, temperature: 22°C or 4°C, blot force: 20, blot time: 4 seconds, wait time: 0 seconds). Grids were clipped and stored in liquid nitrogen until analyzed.

### Cryo-EM

Cryo-EM movies were collected using a Thermo Fisher Scientific Glacios cryo-electron microscope operating at 200 kV equipped with a Gatan K2 Summit direct electron detector at the University of Michigan Life Sciences Institute. Movies were collected at 45,000x magnification (pixel size - 0.980 Å) or 22,000x magnification (pixel size - 2.005 Å) using Leginon software package with exposure time of 8 seconds, frame time of 200 ms, and total dose of 63 e^-^/ Å^2 70^. Movies were collected at defocus of either −3.00 μm or −1.80 μm. Representative still images from movies were obtained using the Leginon software package.

