## Supplemental Information for "*In vitro* synthesis and reconstitution using mammalian cell-free lysates enables the systematic study of the regulation of LINC complex assembly"

* Contributed equally

Corresponding author: Allen P. Liu

**This PDF file includes:**

Supplemental Figures S1 to S5, and Table S1

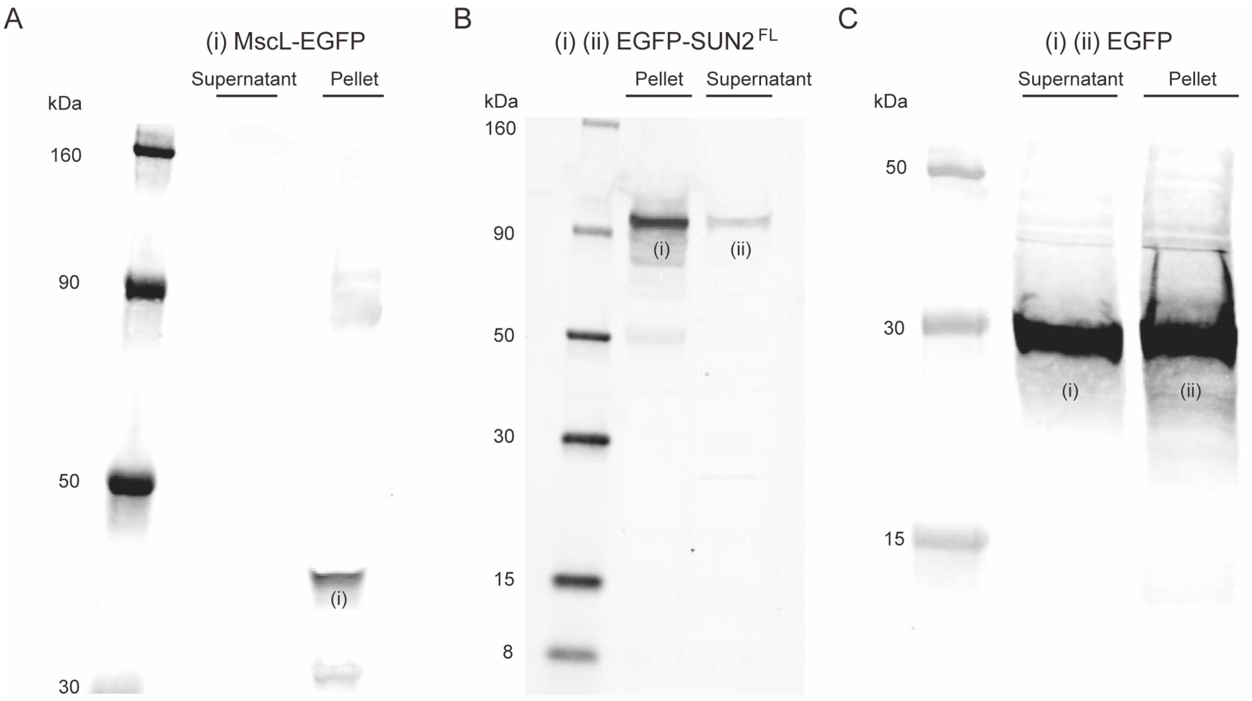


**Figure S1**: **Full in-gel fluorescence images for the data presented in Fig. 1C.**

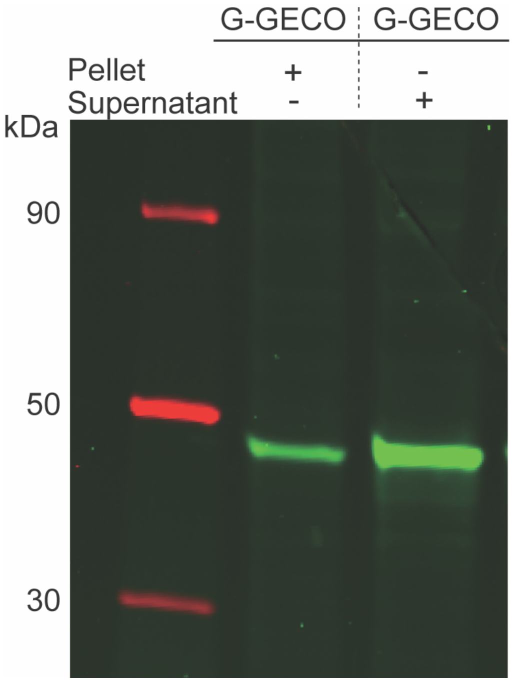


**Figure S2**: **Representative in-gel fluorescence image of the pellet and supernatant fractions of a CFE reaction of the soluble calcium biosensor G-GECO post ultracentrifugation.**

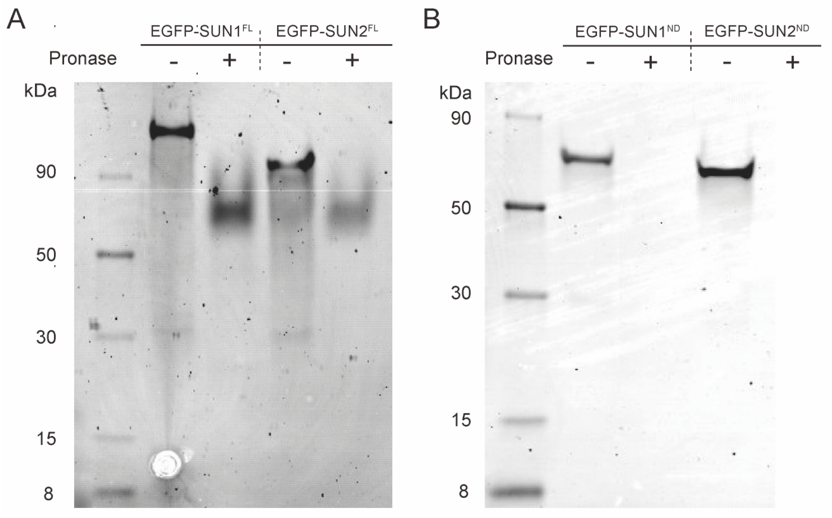


**Figure S3: Full in-gel fluorescence images of the data presented in Fig. 2C.**

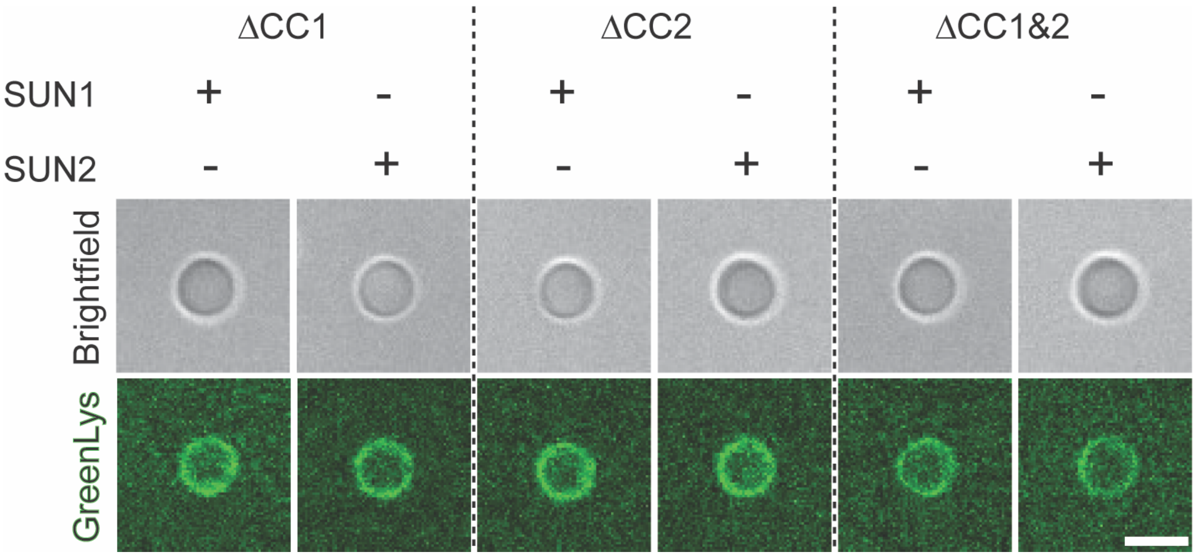


**Figure S4**: **Representative brightfield and confocal fluorescence images of the indicated GreenLys-labeled coiled-coil mutant SUN protein constructs reconstituted in SUPER templates**. Imaging was performed after the SUPER templates were washed twice in 1X PBS containing 6M urea. Scale bar: 5 μm.

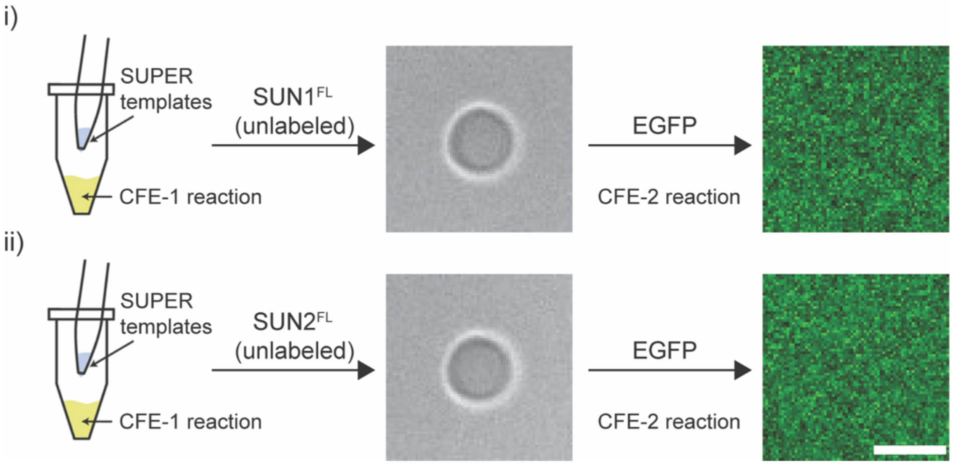


**Figure S5**: **EGFP does not non-specifically associate with SUPER templates exposed to two consecutive rounds of CFE.** i and ii) Left: Schematic of empty SUPER templates being incubated in the first CFE reaction (CFE-1), which synthesized unlabeled FL SUN protein constructs. Middle: Representative brightfield images of SUPER templates containing the indicated reconstituted unlabeled FL SUN protein constructs after being isolated from CFE-1. Right: Representative confocal fluorescence images of the SUPER templates described in the middle panel after being incubated in and isolated from a second CFE reaction (CFE-2) that synthesized soluble EGFP. Scale bar: 5 µm.

| Construct | Forward/Reverse | Primer |
|---|---|---|
| SUN1$^{FL,\Delta CC1}$ | Forward | CTGTGGGTCAAGAATGTGGTTGGAC |
| SUN1$^{FL,\Delta CC1}$ | Reverse | CATGTCACTCTCCTGAGGCCACT |
| SUN1$^{FL,\Delta CC2}$ | Forward | CGCATCCAGGAGACTGTGCAGCT |
| SUN1$^{FL,\Delta CC2}$ | Reverse | GTCATGGTGGAAAGTCATGAAGTCAGTCTTAGC |
| SUN2$^{FL,\Delta CC1}$ | Forward | GAAGTGGGCCTGCTGCCACAG |
| SUN2$^{FL,\Delta CC1}$ | Reverse | GGCCTCCTGGGTCATGCTTTTCC |
| SUN2$^{FL,\Delta CC2}$ | Forward | GCGTCCCTGGGACAGATACTGCAG |
| SUN2$^{FL,\Delta CC2}$ | Reverse | GCGCGCACCCCTGTCTCCAAGAA |
| SUN1$^{FL,\Delta CC1\&2}$ | Forward | CGCATCCAGGAGACTGTGCAGCT |
| SUN1$^{FL,\Delta CC1\&2}$ | Reverse | CATGTCACTCTCCTGAGGCCACT |
| SUN2$^{FL,\Delta CC1\&2}$ | Forward | GCGTCCCTGGGACAGATACTGCAG |
| SUN2$^{FL\ \Delta CC1\&2}$ | Reverse | GGCCTCCTGGGTCATGCTTTTCC |

**Table S1**: Primers for mutagenesis of the mouse SUN1$^{FL}$ (pT7-CFE-SUN1$^{FL}$-His$_6$) and SUN2$^{FL}$ (pT7-CFE-SUN2$^{FL}$-His$_6$) constructs to create the CC-deletion mutants analyzed in this work.
